# Functional Testing of Thousands of Osteoarthritis-Associated Variants for Regulatory Activity

**DOI:** 10.1101/379727

**Authors:** Jason Chesler Klein, Aidan Keith, Sarah J. Rice, Colin Shepherd, Vikram Agarwal, John Loughlin, Jay Shendure

**Author notes:** authors contributed equally (Jason Chesler Klein, Aidan Keith).

## Abstract

To date, genome-wide association studies have implicated at least 35 loci in osteoarthritis, but due to linkage disequilibrium, we have yet to pinpoint the specific variants that underlie these associations, nor the mechanisms by which they contribute to disease risk. Here we functionally tested 1,605 single nucleotide variants associated with osteoarthritis for regulatory activity using a massively parallel reporter assay. We identified six single nucleotide polymorphisms (SNPs) with differential regulatory activity between the major and minor alleles. We show that our most significant hit, rs4730222, drives increased expression of an alternative isoform of *HBP1* in a heterozygote chondrosarcoma cell line, a CRISPR-edited osteosarcoma cell line, and in chondrocytes derived from osteoarthritis patients.

## Main text

Genome-wide association studies (GWAS) have successfully implicated thousands of genetic loci in common human diseases. Most of the underlying signal is believed to derive from variation in non-coding regulatory sequences. However, because of linkage disequilibrium (LD), it has been extraordinarily challenging for the field to identify the variants that causally underlie each association.

Over the past decade, we and others developed massively parallel reporter assays (MPRAs) to increase the throughput at which regulatory sequences can be tested for functional potential^1–4^. An MPRA involves cloning thousands of candidate regulatory sequences to a single reporter gene, transfecting them to a cell line *en masse*, and performing deep sequencing of the resulting transcripts to quantify the degree of transcriptional activation mediated by each candidate regulatory sequence. MPRAs have previously been applied to characterize variants underlying eQTLs (in LCLs)^5^, red blood cell traits (in K562 and human erythroid progenitors/precursors)^6^, cancer-associated common variants (in HEK293 cells)^7^, and adiposity-associated common variants (in HepG2 cells^8^).

Here we sought to apply an MPRA (specifically, STARR-seq^4^) to quantify the relative regulatory potential of SNPs residing on haplotypes implicated in osteoarthritis (OA), with the aim of pinpointing causal variants. We compiled a list of 35 lead SNPs associated with OA in Europeans via GWAS, with minor allele frequencies over 5%^9–26^. Each SNP represents an independent signal with p<5e-8 (genome-wide significant; n=20) or p<5e-5 (genome-wide suggestive; n=15) (**Table S1**). We identified all SNPs in LD with an r^2^>0.8 in Europeans using rAggr (Fig. 1A), resulting in a list of 1,605 candidate SNPs. For the major and minor allele of each SNP, we synthesized 196 nt of genomic sequence, centered on the SNP and flanked by adaptor sequences, on a microarray (230 nt oligos; Fig. 1B).

**Figure 1.**
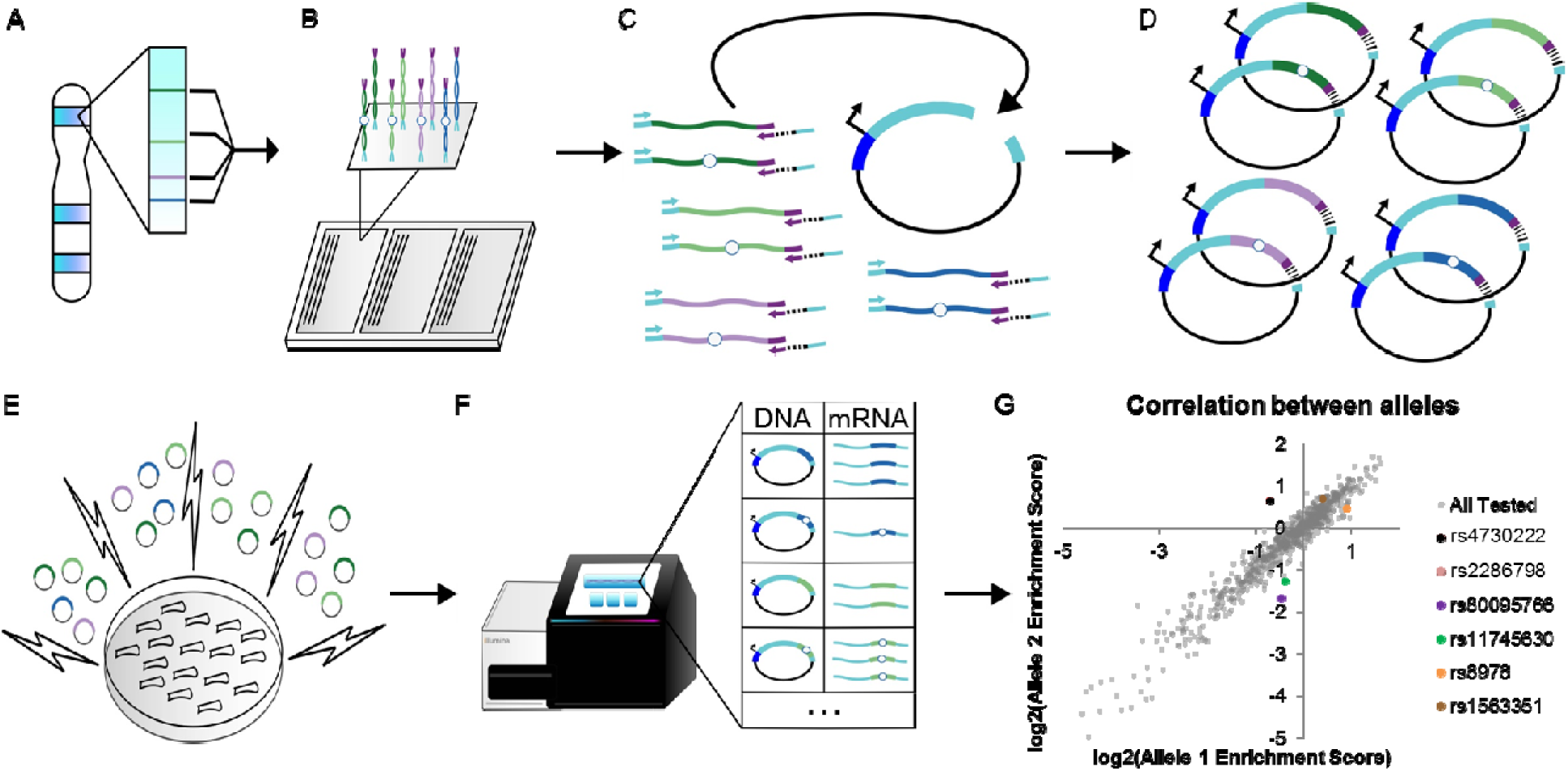
Schematic and results from massively parallel reporter assay. A) For each GWAS-lead SNP, we identified all SNPs in LD with r^2^ > 0.8. Colored lines indicate SNPs in the same LD block. B) For all SNPs, we extracted 196 nt of genomic sequence centered at the SNP, and separately synthesized the minor (hollow circle) and major alleles, flanked by common adaptor sequences (cyan and purple). C-D) We amplified our library from the array via PCR with primers directed at the common adaptors, in the process appending 5 nt degenerate barcode (black lines) and additional sequences homologous to the vector (cyan). We cloned our barcoded library of all major and minor alleles into the STARR-seq vector. Each putative regulatory region is cloned into the 3’ UTR of a reporter gene (cyan) with a minimal promoter (dark blue). E) We transfected our library into Saos-2 cells via electroporation. 48 hours post transfection, we extracted RNA and DNA. F) We determined the abundance of each allele-barcode combination in the mRNA and DNA population through sequencing. G). For each allele, we calculated one activity score as the average log2(RNA/DNA) across all independent measurements. We identified six SNPs with significantly different activity between the major and minor alleles.

During PCR amplification of array-derived oligos, we appended 5 nt degenerate barcodes (Fig. 1C), such that each allele would be represented by multiple independent measurements in the subsequent experiment. We then cloned these barcoded oligos into the human STARR-seq vector and transfected Saos-2 cells, an osteosarcoma cell line. In STARR-seq, the candidate regulatory sequences are located within the transcript itself (Fig. 1D). From the transfected Saos-2 cells, we extracted, amplified and sequenced both DNA and RNA corresponding to the cloned region, and then calculated activity scores as the normalized log2 (ratio of RNA reads / ratio of DNA reads) for each barcode-allele combination (Fig. 1E-F). For all alleles with greater than five independent measurements over three biological replicates (independent transfections of the same library), we averaged the allele activity scores to a single value. Due to bottlenecking during library construction, some alleles were under sampled and therefore excluded from further analysis. Altogether, we obtained activity scores for 1,953 of 3,210 alleles (61%), and activity scores for both alleles of 752 of 1,605 SNPs (47%).

We first asked whether these STARR-seq-based activity scores correlated with biochemical marks for putative enhancers. For this analysis, activity scores corresponding to two alleles of the same SNP were collapsed. We ranked and split the resulting 1,203 activity scores for distinct genomic sequences into quintiles, and then intersected these with datasets of biochemical marks of putative enhancers in cartilage and bone (H3K27ac in bone marrow-derived chondrocytes, H3K27ac in human embryonic limb buds, and ATAC-seq in knee OA cartilage)^27,28^ (Supplementary Fig. 1). The highest scoring quintile was significantly enriched for overlap with H3K27ac ChIP-seq peaks in embryonic limb bud from E41 (2.1-fold, Bonferroni-corrected chi-square p=0.0096), E44 (1.8-fold, p=0.044), and E47 (2.0-fold, p=0.0072), but not with knee OA cartilage ATAC-seq peaks nor bone marrow-derived chondrocyte H3K27ac peaks. These results are in line with our use of an osteosarcoma rather than a cartilage-derived cell line. The highest scoring quintile includes 240 genomic regions, 67 of which overlap putative enhancers from at least one dataset. In contrast, the least active quintile includes 239 genomic regions, only 37 of which overlap putative enhancers from at least one dataset (1.8-fold difference, chi-square p=9.7e-4). Altogether, these enrichments demonstrate that at least a subset of the 1,605 genomic regions tested here correspond to enhancers in OA-relevant tissues. All activity scores are included in **Table S2.**

We next sought to ask whether any alleles are differentially active, focusing on the 752 SNPs for which we successfully measured activity scores for both alleles (**Table S3)**. Overall, activity scores for two alleles of a given SNP were highly correlated, with an overall Spearman correlation of 0.96 (Fig. 1G). This was reassuring, given that each pair of alleles was separately synthesized and cloned, and therefore at non-identical abundances in the STARR-seq library. After correcting for multiple testing with Benjamini-Hochberg (BH) at a 5% FDR, we identified 6 SNPs whose alleles demonstrated significantly differential functional activity in Saos-2 cells (Fig. 1G, **Table S3**). The most significant SNP, rs4730222, is located in the 5’ UTR of several non-canonical isoforms of HMG-Box Transcription Factor 1 (*HBP1).* Other significant SNPs include rs80095766 (intronic to *COG5*), rs2286798 (intronic to *ITIH1*), rs11745630 (downstream of *PIK3R1*), rs6976 (3’ UTR of *GLT8D1*), and rs1563351 (intronic to *LOC102723886*). Two pairs of the six significant SNPs are at the same loci (the ones near *HBP1* and *COG5*, both at chromosome 7q22.3; and the ones near *ITIH1* and *GLT8D1*, both at chromosome 3p21.1).

We chose to further characterize rs4730222 for several reasons. First, this SNP was the most significant hit from our reporter assay, with a substantial difference in activity between the two alleles (2.5-fold increased expression of the minor, disease-associated allele, BH adjusted p-value=2.4e-6). Second, it overlaps several marks for active regulatory elements. Third, we have previously observed reduced expression of the canonical *HBP1* transcript in relevant tissues from carriers of the OA-associated allele^29^, interestingly opposite the effect observed here. HBP1 is a transcriptional repressor that regulates the Wnt-beta-catenin pathway as well as superoxide production, both of which have been implicated in OA development and progression^30,31,32^.

rs4730222 overlaps with several marks associated with active regulatory DNA: H3K27ac (mark for active enhancers and promoters) from bone marrow-derived chondrocytes^28^, human embryonic limb bud at E33, E41, E44 and E47^27^, and ENCODE layered data, H3K4me3 (mark for active promoters) in ENCODE layered data, and ATAC-seq (mark for open chromatin) peaks from articular knee cartilage of OA patients^33^ (Fig. 2A). *HBP1* contains several transcript isoforms, with three probable alternative promoters identified from cap-selected clones and nine validated alternative polyadenylation sites^34^. One of the three probable alternative promoters contains rs4730222 at position +80 relative to the alternative TSS. We therefore hypothesized that the variant may alter expression of this or another *HBP1* isoform. To test this, we first confirmed that the alternative TSS overlapping rs4730222 is utilized in Saos2 and SW1353 cells (a chondrosarcoma cell line) with qRT-PCR with primers contained within this 5’ UTR as well as spanning to the following exon. However, despite multiple attempts, we failed to successfully amplify the isoform from the alternative TSS to the canonical stop, suggesting that the alternative TSS may belong to a truncated isoform of the *HBP1* transcript. Based on PCR and Sanger sequencing, the truncated isoform is most likely ENST00000497535, which contains the alternative 5’ UTR and first two exons (Fig. 2A, Supplementary Fig. 2).

**Figure 2.**
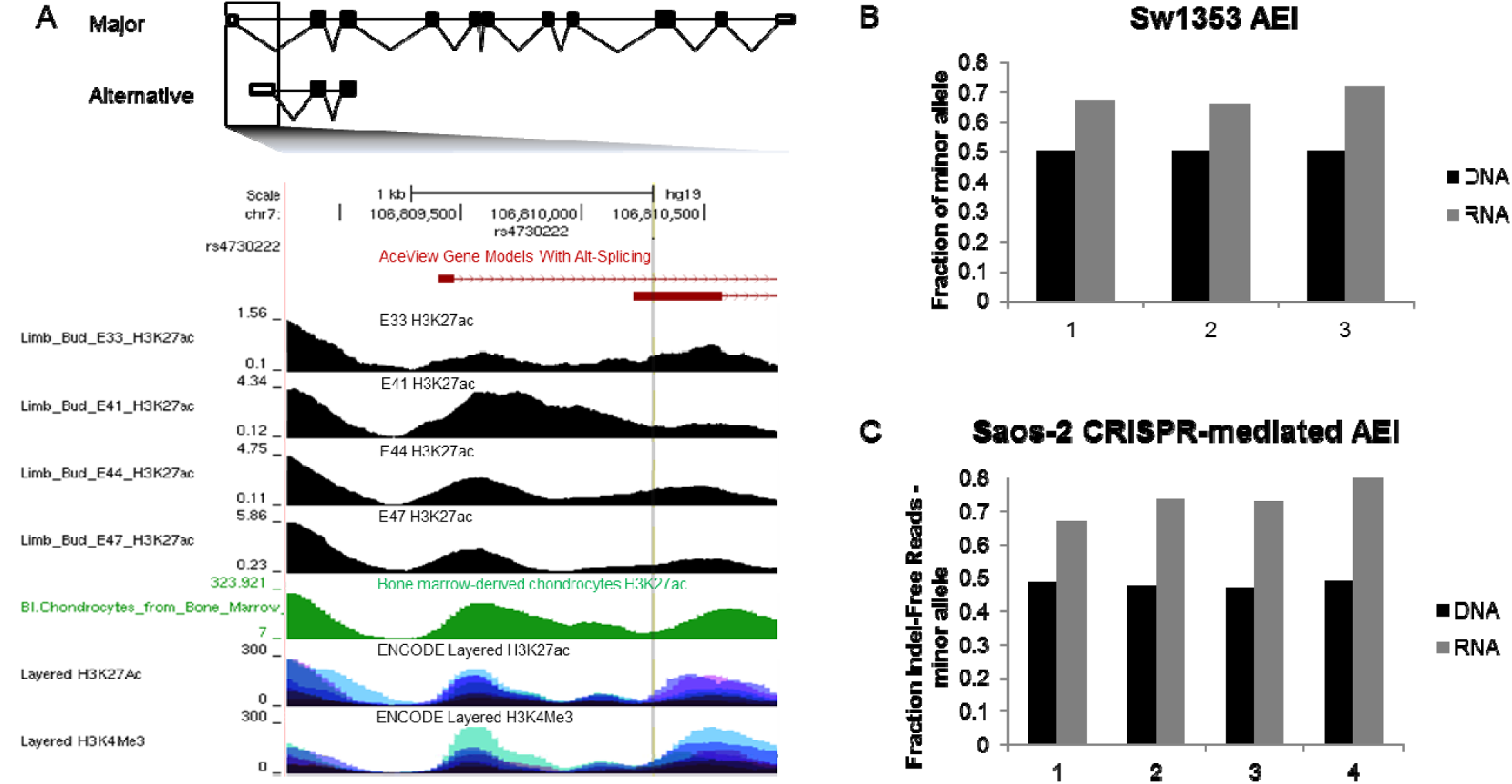
Functional Validation of rs4730222. A) Gene model of *HBP1* ensemble isoform ENST00000468410 (major) and ENST00000497535 (alternative). UCSC genome browser zoomed in to the two transcriptional start sites (http://genome.ucsc.edu). Rs4730222, within th 5’ UTR of the alternative transcript, is indicated by a vertical grey line traversing the annotation tracks. The four black tracks are H3K27ac performed in human embryonic limb bud at E33, E41, E44 and E47 respectively^27^. The green track is H3K27ac data from chondrocytes derived from cultured bone marrow mesenchymal stem cells^28^. Layered H3K27ac is H3K27ac ChIP-seq (a marker for active enhancers and active promoters) layered from GM12878, H1-hESC, HSMM, HUVEC, K562, NHEK, and NHLF cells. Layered H3K4me3 is H3K4me3 ChIP-seq (marker for active promoters) layered from the same seven ENCODE cell lines. B) Allelic expression imbalance in SW1353, a chondrosarcoma cell line heterozygote for rs4730222. Black bar represent the fraction of the minor allele in DNA and grey bars indicate the fraction of the minor allele in cDNA. C) Allelic expression imbalance in Saos-2 cells with the minor allele of rs4730222 introduced through CRISPR-mediated HDR. Bars are the same as in Fig. 2B.

After confirming that rs4730222 is transcribed as part of a *HBP1* isoform in osteogenic and chondrogenic cells (albeit not to the canonical stop), we genotyped rs4730222 in several cell lines (SW1353, Tc28a/2, Saos-2, chondrogenic progenitor cells) in hopes of identifying a heterozygous line. As SW1353 was heterozygous for rs4730222, we tested it for allelic expression imbalance (AEI) of the transcribed SNP. In all three biological replicates, we observed a significant allelic imbalance (Fisher’s exact test, p<1e-5), with the minor allele showing a 1.3 to 1.4-fold relative enrichment in RNA/DNA compared to the major allele. This was more modest but directionally concordant with 2.5-fold enrichment of the minor allele in the reporter assay (Fig. 2B).

The OA-implicated haplotype at 7q22.3 is a ~500 kb region of high linkage disequilibrium, and consequently accounts for 283 of the 1,605 SNPs tested here. The observed AEI of the *HBP1* alternative isoform in SW1353 cells, which are likely heterozygous for the entire OA-associated haplotype, could be consequent to any or several of these SNPs, or to other forms of genetic variation. To test whether rs4730222 causally underlies the allelic imbalance of the *HBP1* alternative transcript, we introduced the minor allele of rs4730222 into Saos-2 cells, which are homozygous for the major allele, through CRISPR-mediated homology directed repair (HDR). We generated four biological replicates (*i.e.* independently edited cell populations), and quantified the RNA-DNA ratio of indel-free reads derived from the major vs. minor allele. Similar to the SW1353 AEI (1.3 to 1.4-fold), we identified a 1.4 to 1.6-fold relative enrichment in the RNA/DNA ratio for the minor allele compared to the major allele (Fisher’s exact test, p<1e-5; Fig. 2C). This confirms that the minor allele at rs4730222 causally underlies upregulation of the *HBP1* alternative isoform.

We next sought to test whether rs4730222 drives increased transcription of the alternative TSS in osteoarthritic tissue. To do so, we tested for AEI in chondrocytes derived from OA patients. For each of nine patients heterozygous for rs4730222, we extracted RNA and DNA and amplified and sequenced the 5’UTR containing the SNP (three technical replicates per patient) from DNA and cDNA. Despite low cDNA concentrations and low expression of the alternative TSS, we observed an overall trend in AEI in accordance with our reporter assay and cell models (median AEI=1.12, Mann Whitney U-test on 27 DNA ratios vs. 27 RNA ratios, p=0.003). Interestingly, one patient appeared to exhibit AEI in the opposite direction^35,36^ (Supplementary Fig. 3).

In summary, we set out to functionally test the regulatory effects of 1,605 SNPs that potentially underlie 35 GWAS signals for osteoarthritis. We succeeded in generating reproducible measurements of regulatory activity for about two-thirds of the regions tested, and of differential regulatory activity for about half of the regions tested. The most highly active regions in our assay were ~2-fold enriched for biochemical marks associated with enhancers. We furthermore identified six SNPs, which each drove differential expression at an FDR of 5%. The most significant of these, rs4730222, resides in the 5’ UTR of multiple isoforms of *HBP1*, a transcriptional repressor. The minor, OA-associated risk allele of rs4730222 increases transcription of alternative isoform(s) of *HBP1*. We validated this finding in SW1353 cells, CRISPR-edited Saos2 cells, and in chondrocytes derived from knee OA patients.

We previously described reduced expression of the canonical *HBP1* transcript in OA tissue, relative to healthy tissue^29^. Here, we do not observe AEI of the canonical *HBP1* transcript, but rather AEI of an alternative transcript. We speculate that the impact of rs4730222, wherein the disease-associated allele consistently increases expression of an alternative transcript, secondarily reduces expression of canonical *HBP1*. One possibility is that the alternative isoform encodes a non-coding RNA with an upstream open reading frame (uORF). This isoform of *HBP1* may have a *trans* effect on the canonical transcript, or have its own, yet uncharacterized, function in the cell. Additionally, alternative TSSs have been shown to modulate translational efficiency and tissue specificity of genes^37,38^. Through one or several of these mechanisms, the expression levels of this isoform might modulate HBP1 levels in certain tissues. Finally, we note that the rs4730222-overlapping isoform expressed in SW1353 and Saos2 is a truncated version of *HBP1*. Therefore, it may disrupt endogenous activity of the gene, *e.g.* by acting as a dominant-negative. Distinguishing between these mechanistic possibilities for rs4730222 and *HBP1,* as well as for other MPRA-prioritized OA-associated SNPs, should be a high priority for the field.

## Acknowledgements

The authors thank the Shendure and Loughlin labs, particularly J. Alexander, M. Gasperini, and S. Kim, for helpful discussions. This work was funded by grants from the National Institutes of Health (UM1HG009408, R01CA197139, R01HG006768) to J.S. J.C.K. was supported in part by 1F30HG009479 from the NHGRI. J.S. is an investigator of the Howard Hughes Medical Institute. S.J.R., C.S. and J.L. were supported by Arthritis Research UK (grant 20771), and by the Medical Research Council and Arthritis Research UK as part of the MRC-Arthritis Research UK Centre for Integrated Research into Musculoskeletal Aging (CIMA, grant references JXR 10641, MR/P020941/1 and MR/R502182/1).

## Author Contributions

The project was conceived and designed by J.C.K., A.K., J.S., and J.L. S.J.R. and C.S. collated and generated the OA patient DNA and cDNA samples. J.L. compiled OA lead variants. J.C.K. and A.K. performed all experiments and analyses. V.A. performed additional analysis and modeling. J.C.K., A.K. and J.S. wrote the manuscript. All authors read and approved the final version of the manuscript.

## Supplementary Figures

**Supplementary Table 1.**
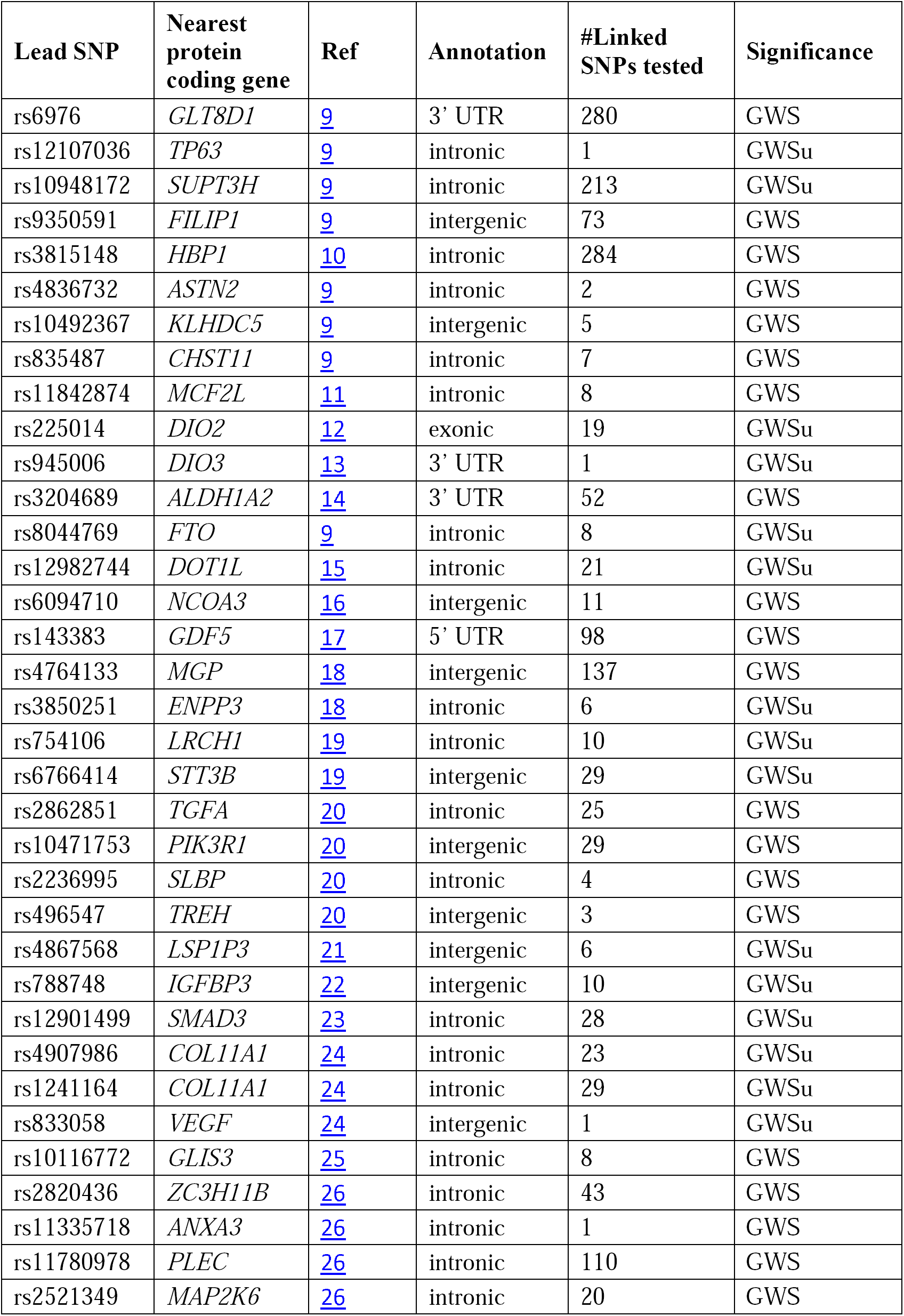
Lead SNPs. List of 20 genome-wide significant (GWS) and 15 genome-wide suggestive (GWSu) variants compiled in May 2017.

**Supplementary Figure 1.**
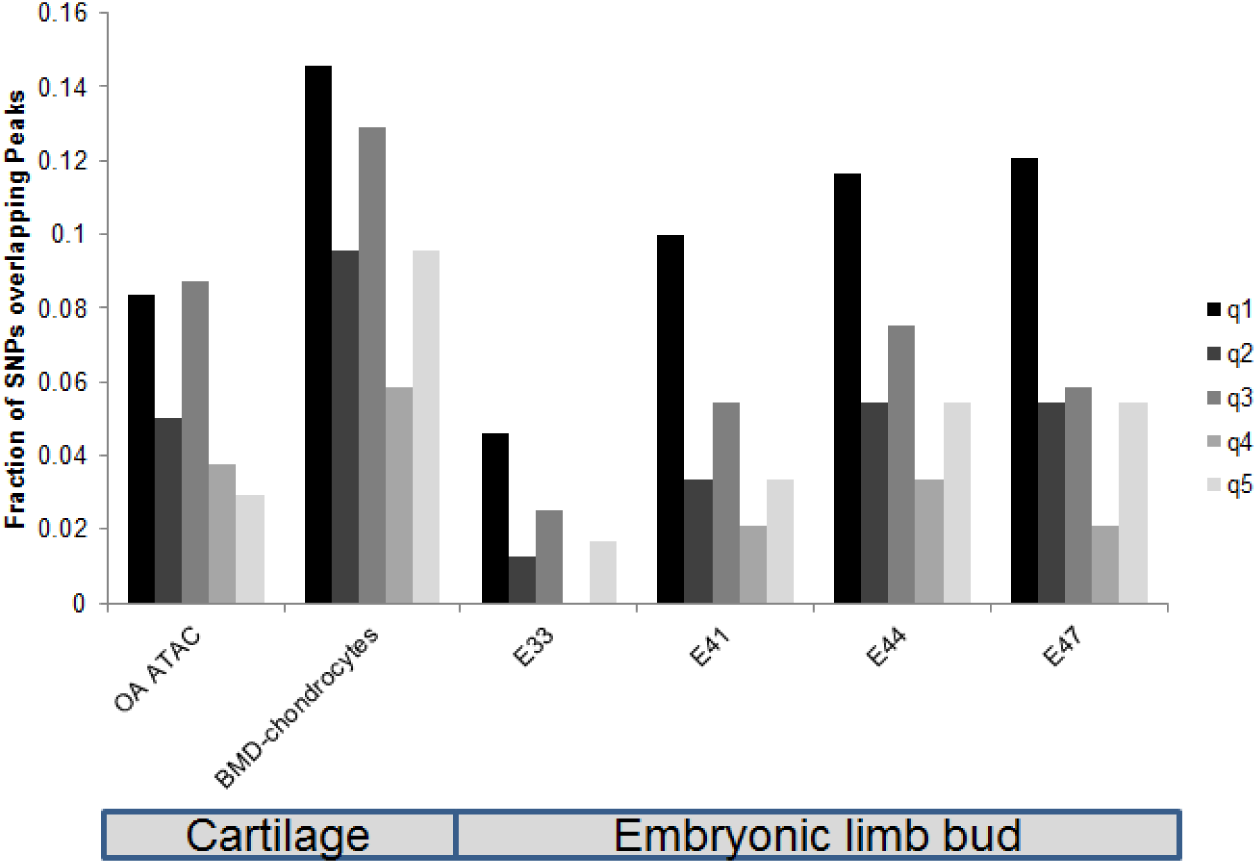
Overlap between tested sequences and enhancer marks. The 1,203 SNPs were split into 5 quintiles of ~240 sequences each, based on their normalized RNA/DNA activity score. Q1 refers to the sequences with the highest activity scores and q5 refers to the sequences with the lowest activity scores. We then overlapped each quintile with peaks called from OA ATAC-seq^33^, BMD-chondrocytes H3K27ac ChIP-seq^28^, and human embryonic limb bud H3K27ac ChIP-seq from E33, E41, E44, and E45^27^. Y-axis is the fraction of the 240 SNPs in each quintile overlapping peaks.

**Supplementary Figure 2.**
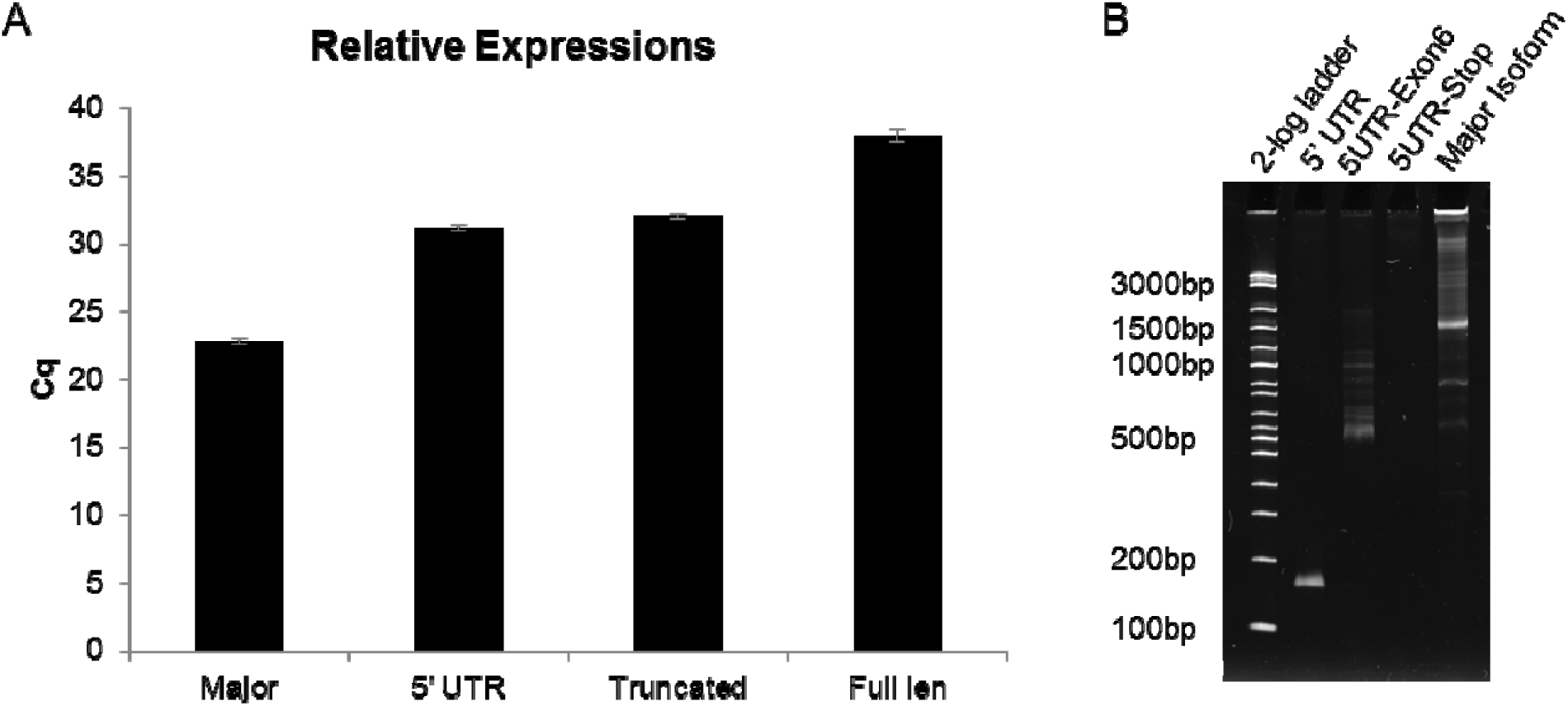
Relative expression of different HBP1 isoforms. A) Y axis is the Cq value from RT-qPCR from SW1353 cells. The major allele uses a forward primer at the start codon and reverse primer at the conserved stop codon of the major isoforms. The 5’ UTR primer set amplifies a short product, entirely within the alternative 5’ UTR. The truncated primer set amplifies both ENST00000497535 and ESNT00000485846. The full length primer set includes a forward primer in the alternative TSS and reverse primer at the stop codon of the major isoform. We do not identify any full length product utilizing the alternative TSS. B) Gel of qPCR products. First lane is a 2-log ladder. Second lane is the 5’ UTR amplification. Expected size is 156 bp. Third lane is the truncated amplification. The primer set should amplify both ENST00000497535 (expected size 548bp) and ENST00000485846 (expected size 846bp). However, ENST00000485846 contains an internal exon while ESNT00000497535 does not. We ran the PCR product on a gel and Sanger sequenced the purified PCR product, which did not include the internal exon. Fourth lane is amplifying from the alternative 5’ UTR to canonical stop. There was no amplification product. Fifth lane is the major isoform (amplifying from canonical start to canonical stop. Expected size is 1,455 bp.

**Supplementary Figure 3.**
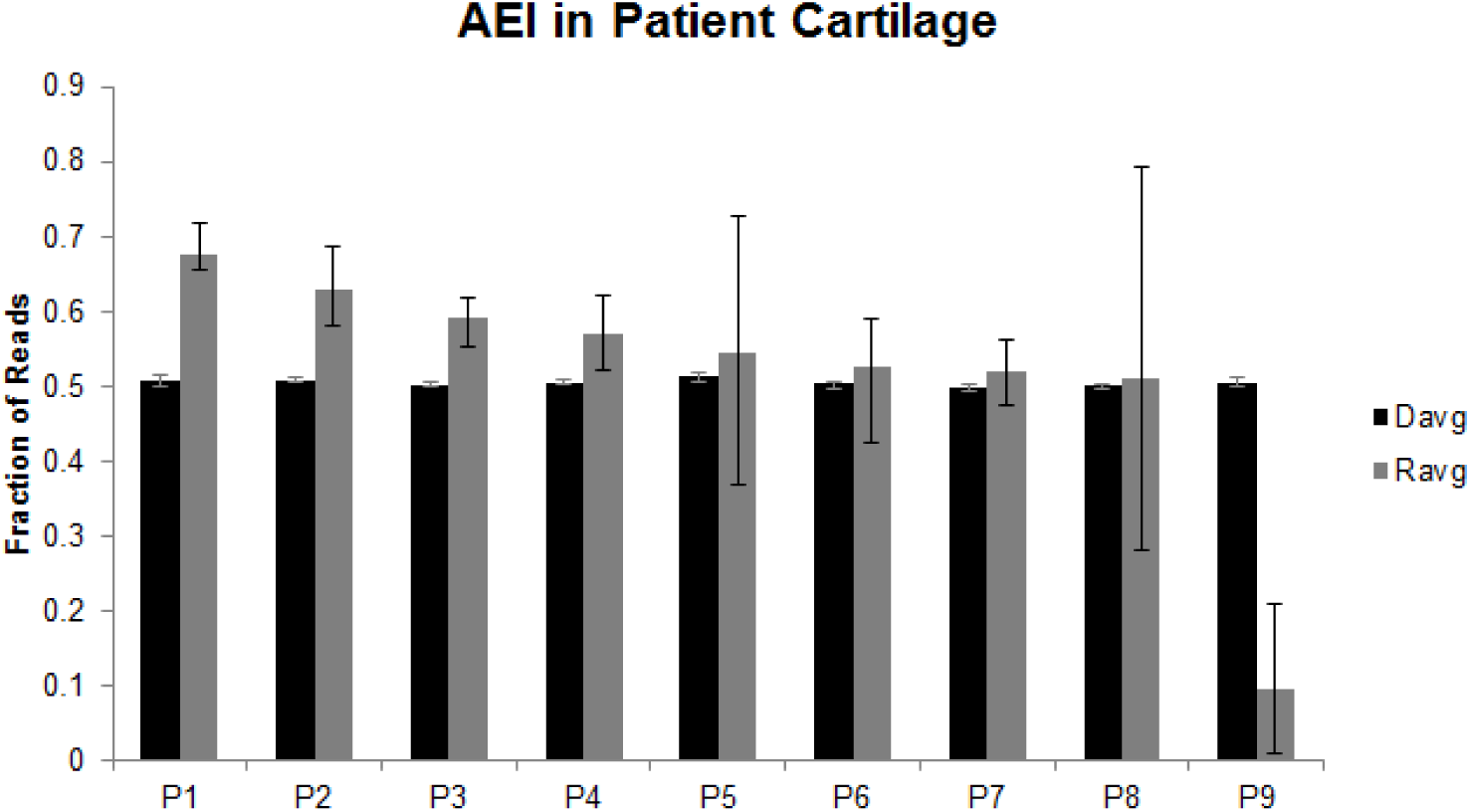
Allelic Expression Imbalance (AEI) in cartilage from patients receiving total knee replacements. Black bars are the fraction of DNA reads aligning to the minor allele. Grey bars are the fraction of RNA reads aligning to the minor allele. Bars indicate th minimum and maximum fraction from three technical replicates.

## Methods

### Identification and design of target SNPs

We selected SNPs that had a minor allele frequency >5% and had been reported as being associated with OA in European populations at a significance level that surpassed or approached the genome-wide threshold of <5e-8. The deadline date for inclusion was May 2017. In total, 35 SNPs were identified, each representing an independent association signal (Table S1). We ran rAGGr on our list of 35 candidate SNPs to identify all variants with a minimum minor allele frequency >= 0.001 in linkage disequilibrium with an r^2^>0.8 in Europeans (CEU+FIN+GBR+IBS+TSI) based on 1000 Genomes, Phase 3, Oct 2014. We then filtered out any polymorphisms greater than one nucleotide, resulting in a list of 1,605 SNPs. For each variant, we extracted 196 nt of genomic sequence centered around the SNP using BEDTOOLS getfasta, and edited the SNP to create both the minor and major alleles (3,210 sequences). To each 196 nt sequence, we appended HSS_clon_F (5’ - TCTAGAGCATGCACCGG - 3’) to the 5’ end and DO_R6 (5’-GCCGGTCAGAATGATGG -3’) to the 3’ end. We then ordered the 3,210 sequences in duplicate as part of an Agilent 244K 230-mer array.

### Library Generation

We amplified our sequences off of the Agilent array with HSS_clon_F and R6_5N_HSSR (5’-CCGGCCGAATTCGTCGANNNNNCCATCATTCTGACCGGC -3’) using KAPA HiFi HotStart ReadyMix in a 50 uL reaction with 0.75 ng DNA and SYBR Green on a MiniOpticon Real-Time PCR system (Bio-Rad) and stopped the reaction before plateauing (13 cycles). This reaction amplified our library, added a 5 nt degenerate barcode to each sequence, and added both adapters for cloning into the human STARR-seq vector. We purified the PCR product using a 1.5x AMPure cleanup following manufacturer’s protocol. We then ligated 6 ng of our purified PCR into 25 ng of linearized human STARR-seq backbone using the NEBuilder HiFi DNA Assembly Cloning Kit following manufacturer’s protocol. We transformed 1.2 uL of the ligation product in 50 uL of NEB C3020 cells, grew up overnight in 100 mL of LB+Amp, and extracted the library using a Zymo Research ZymoPURE Plasmid Midiprep Kit.

### STARR-seq Screen

We transfected 1.5 million Saos-2 cells with 20 ug of our library in triplicate using the Thermo Fisher Scientific Neon Transfection System with resuspension buffer R at 1250V, 40ms, 1 shock, with 100 uL pipettes, in triplicate. After electroporation, we added the cells to 10cm plates with pre-warmed media (McCoy’s 5A with 10% FBS and 1x Pen/Strep). 48 hours post transfection, we extracted both DNA and RNA from each replicate using the Qiagen ALLPrep DNA/RNA Mini Kit. DNA was eluted in 80 uL and RNA was eluted in 30 uL. RNA was treated with Thermo Fisher Scientific TURBO DNase following manufacturer’s protocol and reverse transcribed using Thermo Fisher Scientific SuperScript III Reverse Transcriptase in a 20 uL reaction with 8 uL of RNA. For each replicate, we amplified DNA in two reactions, each with 2 ug of DNA using NEBNext High Fidelity 2X PCR Master Mix with primers HSS_NF_pu1 (5’-CTAAATGGCTGTGAGAGAGCTCAGGTACAACTGATCTAGAGCATGCACC -3’) and HSS_R_pu1.(5’-ACTTTATCAATCTCGCTCCAAACCCTTATCATGTCTGCTCGAAGC -3’) and stopped before plateauing (15 cycles). After PCR, products were purified with a 1.5x AMPure cleanup, and pooled together. For each replicate, we also amplified cDNA in two reactions, each with 10 uL of RT product in 50 uL reactions with NEBNext High Fidelity 2X PCR Master Mix with primers HSS_F_pu1 (5’-CTAAATGGCTGTGAGAGAGCTCAGGGGCCAGCTGTTGGGGTGTCCAC-3’) and HSS_R_pu1 (5’-ACTTTATCAATCTCGCTCCAAACCCTTATCATGTCTGCTCGAAGC -3’) and stopped before plateauing (18-20 cycles). After PCR, products were purified with a 1.5x AMPure cleanup, eluted in 50 uL each, and pooled together. For the cDNA samples, we performed a nested reaction using KAPA HiFi HotStart ReadyMix in a 50 uL reaction with 1 uL of the pooled outer PCR reaction with HSS-NF-pu1 and pu1R (5’-ACTTTATCAATCTCGCTCCAAACC -’3) and stopped before plateauing (7 cycles). Reactions were purified with a 1.5x AMPure cleanup and eluted in 50 uL each. Flow cell adapters and indexes were added to all DNA and cDNA reactions through an additional round of PCR using Kapa HiFi HotStart ReadyMix in 50 uL reactions with 1 uL of the first DNA PCR or 1 uL of the inner cDNA PCR using an indexed pu1_P5 primer (5’-AATGATACGGCGACCACCGAGATCTACACNNNNNNNNNNACGTAGGCCTAAATGGCTGTGAGAGAGCTCAG -3’) and an indexed pu1_P7 primer (5’-CAAGCAGAAGACGGCATACGAGATNNNNNNNNNGACCGTCGGCACTTTATCAATCTCGCTCCAAACC -3’) and stopped before plateauing (6 cycles). The libraries were sequenced on an Illumina NextSeq 500/550 v2 300 cycle mid-output kit.

### Analysis of STARR-seq Screen

We aligned all sequencing reads to a reference fasta file of our variants using BWA mem and extracted reads from error-free molecules^39^. Each variant contained several different 5 nt barcodes added through PCR. We counted the number of reads from each replicate for each variant-barcode combination in the DNA and cDNA pool. If there were at least 10 DNA reads and at least 1 RNA read, we calculated an activity score as the log2(number of RNA reads from the variant-barcode combination normalized to the total number of RNA reads, divided by the number of DNA reads from the variant-barcode combination normalized to the total number of DNA reads). We then combined all variant-barcode activity scores from each replicate, and for any variant with at least five different measurements, we averaged the activity score for a final activity score for each variant. This resulted in activity scores for 1,953 of the 3,210 alleles. 752 of the 1,605 variants contained measurements for both alleles. For each of the 752 variants with measurements for both alleles, we tested whether the 2 alleles drove different expression by performing a Mann-Whitney U Test for each variant using SciPy v0.19.1 with Python v2.7.3. We then performed a Benjamini-Hochberg correction with an FDR = 0.05 to correct for multiple testing.

### Allelic Imbalance of rs4730222 in SW1353 cells

We first genotyped several osteogenic and chondrogenic cell lines for rs4730222 (SW1353, Tc28a/2, Saos-2, chondrogenic progenitor cells) using rs4730222_sangerF (5’-TACGCAGTTCGAATGAATGGGCTC -3’) and rs4730222_sangerR (5’-AGCTACAAAAACCTGGCTGTCCAC -3’). PCR products were purified with a 1.5x AMPure cleanup and Sanger sequenced with rs4730222_sangerF.

We then tested for allelic imbalance of rs4730222 in the isoforms expressing the SNP in SW1353. We performed three independent DNA and RNA extractions using the Qiagen ALLPrep DNA/RNA Mini Kit. DNA was eluted in 80 uL and RNA was eluted in 30 uL. RNA was treated with TURBO DNase and reverse transcribed with SuperScript III Reverse Transcriptase. We then amplified the 5’UTR around rs4730222 from each DNA and cDNA sample using KAPA HiFi HotStart ReadyMix in a 50 uL reaction with 100 ng DNA or 5 uL cDNA with HBP1_5UTR_F_pu1 (5’-CTAAATGGCTGTGAGAGAGCTCAGAGTCCGGGCTGCGGTCACATGATG -3’) and HBP1_5UTR_R_pu1 (5’-ACTTTATCAATCTCGCTCCAAACCAGCTACAAAAACCTGGCTGTCCAC -3’) and stopped DNA reactions at 25 cycles and cDNA reactions at 32 cycles. Products were purified using a 1.5x AMPure cleanup, and flow cell adapters and indexes were added using an indexed pu1_P5 primer and an indexed pu1_P7 primer. Libraries were spiked into a Miseq v2 300 cycle run. Reads were aligned to a fasta reference file using BWA mem and the number of perfect reads coming from both alleles was quantified from both DNA and cDNA from each replicate.

### CRISPR Knock-in of rs4730222 in Saos-2 cells

rs4730222 falls within a potential Cas9 PAM site (5’-ACGCGATGAATGGCGAAAGA G**G**G - 3’). We therefore designed a guideRNA that would target rs4730222, so that the minor-allele donor would not be re-cut. We ordered the following oligos from IDT: rs4730222_guideF (5’-CACCGACGCGATGAATGGCGAAAGA -3’) and rs4730222_guideR (5’-AAACTCTTTCGCCATTCATCGCGTC -3’) and followed the Zhang lab protocol to clone them into the px458 plasmid (SpCas9-2A-EGFP and single guide RNA).

We created our donor vector in two steps. First, we amplified a 1,459 bp region around rs4730222 with the following primers, which also append 16 bp homologous sequence to puc19 onto each side of the amplicon: HBP1_puc19F (5’-TCGGTACCCGGGGATCAAGTAGGAAAGTTTCGGTTGAGGAG -3’) and HBP1_puc19R (5’-TCGACTCTAGAGGATCAACTGAACAGATGACCGACTCTACC -3). We then cloned this into a linearized puc19 plasmid using Clontech’s In-Fusion HD Cloning Kit following manufacturer’s protocol, transformed into Stellar Competent cells, grew up a single colony and extracted plasmid using the Zymo Research ZymoPURE Plasmid Midiprep Kit. We then re-linearized the puc19-HBP1 wild-type plasmid via PCR with puc19_HBP1-linF (5’-GTGGGGGATGGACTTGGCGTG -3’) and puc19-HBP1-linR (5’-CTCCTCAACCGAAACTTTCCTACTT -3’). We also amplified a small region around rs4730222, while mutating the SNP, using mut_insF (5’-AAGTAGGAAAGTTTCGGTTGAGGAG -3’) and mut_insR (5’-CCAAGTCCATCCCCCACGCTCTTTCGCCATTCATCGCG -3’). We then cloned the mutated insert into puc19-HBP1-wt using the In-Fusion HD Cloning Kit and grew up a single colony with the minor allele at rs4730222 flanked by 600-850 bp of homology on each side.

We transfected 1 million Saos-2 cells with 10 ug of our px458-rs4730222 guide and 10 ug of our donor library containing the minor allele using the Neon Transfection system as described above. 72 hours post transfection, we performed FACS on a BD FACS Aria III to isolate ~150,000 GFP+ cells (transfected with px458), which we then expanded. On day 10 post transfection, we extracted DNA and RNA, performed reverse transcription with Superscript III, and amplified the region surrounding rs4730222 from both DNA (using HBP1_5UTR_F_pu1 and HBP_DNA_Routside (5’-TAGGTGGGCAATCCTGGGAGAAGGTAC -3’)), and RNA (using HBP1_5UTR_F_pu1 and HBP_RNA_Routside (5’-TGCCAGATTCTGACTCACTATTTGC -3’)) in 50 uL reactions using KAPA HiFi 2x ReadyMix. We then purified the PCR reactions with a 1.5x AMPure cleanup, eluted in 50 uL, and used 1 uL in a nested reaction with pu1L (5’-CTAAATGGCTGTGAGAGAGCTCAG -3’) and HBP1_5UTR_R_pu1. Reactions were purified with a 1.5x AMPure cleanup, and flow cell adapters and indexes were added using an indexed pu1_P5 primer and an indexed pu1_P7 primer. Libraries were spiked into a Miseq v2 300 cycle run. Reads were aligned to a fasta reference file using BWA mem and the number of perfect reads coming from both alleles was quantified from both DNA and cDNA from each replicate.

### Allelic imbalance of rs4730222 in osteoarthritis patients’ chondrocytes

Cartilage tissue samples were obtained from OA patients who had undergone joint replacement surgery at the Newcastle upon Tyne NHS Foundation Trust hospitals. The Newcastle and North Tyneside Research Ethics Committee granted ethical approval for the collection, with each donor providing verbal and written informed consent (REC reference number 14/NE/1212). Our patient ascertainment criterion has been described in detail previously^40,41^. The cartilage was removed from the joint using a scalpel and was collected distal to the OA lesion. The tissue samples were stored frozen at −80°C and ground to a powder using a Retsch Mixermill 200 (Retsch Limited) under liquid nitrogen. Nucleic acids were then extracted from the ground tissue using TRIzol reagent (Life Technologies) according to the manufacturer’s instructions, with the upper aqueous phase separated for RNA isolation, while the interphase and lower organic phase were used to isolate DNA. RNA was reverse transcribed using the SuperScript First-Strand cDNA synthesis kit (Invitrogen). Matched DNA and cDNA were amplified with KAPA HiFi 2x ReadyMix and SYBR Green and halted before plateauing. The primers sequences were as follows: 5’-CTAAATGGCTGTGAGAGAGCTCAGAGTCCGGGCTGCGGTCACATGATG-3’; and 5’-ACTTTATCAATCTCGCTCCAAACCAGCTACAAAAACCTGGCTGTCCAC-3’.

All samples were purified with a 1.5x AMPure cleanup following manufacturer’s instructions and eluted in 50 uL Qiagen Elution Buffer. 1 uL of purified product was then indexed for Illumina sequencing using an indexed pu1_P5 primer and an indexed pu1_P7 primer. Libraries were spiked into a Miseq v2 300 cycle run. Reads were aligned to a fasta reference file using BWA mem and the number of aligning reads coming from both alleles was quantified from both DNA and cDNA from each replicate. For statistical analysis, a Mann Whitney U-test was performed comparing DNA vs. RNA abundances of the minor allele for the 54 values (27 for DNA and 27 for RNA; nine patients x three replicates).

### Characterization of HBP1 rs4730222-containing isoforms

We designed the following set of primers to differentiate between different HBP1 isoforms: Major (1stExonF: 5’-GTGTGGGAAGTGAAGACAAATCAGATGC -3’ and LastExonR: 5’-CTTCCACCTGTCACCAAGGATCACAC -3’), 5’ UTR (UTR_qPCR_F: 5’-CAGTCTCCGCCTTTCAACCTATG -3’ and UTR_qPCR_R: 5’-ATGAACTCGAGTGTAGAGTGCACAG -3’), Truncated (UTR_qPCR_F and Exon6_R: CCACCTCATTTTCACGGTAAGTAG -3’) and Full Len (UTR_qPCR_F and LastExonR). We performed technical triplicates for each qPCR using KAPA Robust 2x Hotstart Readymix with cDNA from wild-type SW1353 cells, letting the reaction go for 40 cycles. We then ran products on a gel, and differentiated between ENST00000497535 and ENST00000485846 based both on size and Sanger sequencing.

## REFERENCES

1. Patwardhan, R. P. et al. High-resolution analysis of DNA regulatory elements by synthetic saturation mutagenesis. Nat. Biotechnol. 27, 1173–1175 (2009).

2. Patwardhan, R. P. et al. Massively parallel functional dissection of mammalian enhancers in vivo. Nat. Biotechnol. 30, 265–270 (2012).

3. Melnikov, A. et al. Systematic dissection and optimization of inducible enhancers in human cells using a massively parallel reporter assay. Nat. Biotechnol. 30, 271 (2012).

4. Arnold, C. D. et al. Genome-wide quantitative enhancer activity maps identified by STARR-seq. Science 339, 1074–1077 (2013).

5. Tewhey, R. et al. Direct Identification of Hundreds of Expression-Modulating Variants using a Multiplexed Reporter Assay. Cell 172, 1132–1134 (2018).

6. Ulirsch, J. C. et al. Systematic Functional Dissection of Common Genetic Variation Affecting Red Blood Cell Traits. Cell 165, 1530–1545 (2016).

7. Liu, S. et al. Systematic identification of regulatory variants associated with cancer risk. Genome Biol. 18, 194 (2017).

8. Vockley, C. M. et al. Massively parallel quantification of the regulatory effects of noncoding genetic variation in a human cohort. Genome Res. 25, 1206–1214 (2015).

9. Consortium, A. & Others. arcOGEN Collaborators, Zeggini E, Panoutsopoulou K, Southam L, Rayner NW, et al. Identification of new susceptibility loci for osteoarthritis (arcOGEN): a genome-wide association study. Lancet 380, 815–823 (2012).

10. Kerkhof, H. J. M. et al. A genome-wide association study identifies an osteoarthritis susceptibility locus on chromosome 7q22. Arthritis Rheum. 62, 499–510 (2010).

11. Day-Williams, A. G. et al. A variant in MCF2L is associated with osteoarthritis. Am. J. Hum. Genet. 89, 446–450 (2011).

12. Meulenbelt, I. et al. Identification of DIO2 as a new susceptibility locus for symptomatic osteoarthritis. Hum. Mol. Genet. 17, 1867–1875 (2008).

13. Meulenbelt, I. et al. Meta-analyses of genes modulating intracellular T3 bio-availability reveal a possible role for the DIO3 gene in osteoarthritis susceptibility. Ann. Rheum. Dis. 70, 164–167 (2011).

14. Styrkarsdottir, U. et al. Severe osteoarthritis of the hand associates with common variants within the ALDH1A2 gene and with rare variants at 1p31. Nat. Genet. 46, 498–502 (2014).

15. Evangelou, E. et al. The DOT1L rs12982744 polymorphism is associated with osteoarthritis of the hip with genome-wide statistical significance in males. Ann. Rheum. Dis. 72, 1264–1265 (2013).

16. Evangelou, E. et al. A meta-analysis of genome-wide association studies identifies novel variants associated with osteoarthritis of the hip. Ann. Rheum. Dis. 73, 2130–2136 (2014).

17. Miyamoto, Y. et al. A functional polymorphism in the 5’ UTR of GDF5 is associated with susceptibility to osteoarthritis. Nat. Genet. 39, 529–533 (2007).

18. den Hollander, W. et al. Genome-wide association and functional studies identify a role for matrix Gla protein in osteoarthritis of the hand. Ann. Rheum. Dis. 76, 2046–2053 (2017).

19. Panoutsopoulou, K. et al. Radiographic endophenotyping in hip osteoarthritis improves the precision of genetic association analysis. Ann. Rheum. Dis. 76, 1199–1206 (2017).

20. Castaño-Betancourt, M. C. et al. Novel Genetic Variants for Cartilage Thickness and Hip Osteoarthritis. PLoS Genet. 12, e1006260 (2016).

21. Yau, M. S. et al. Genome-Wide Association Study of Radiographic Knee Osteoarthritis in North American Caucasians. Arthritis Rheumatol 69, 343–351 (2017).

22. Evans, D. S. et al. Genome-wide association and functional studies identify a role for IGFBP3 in hip osteoarthritis. Ann. Rheum. Dis. 74, 1861–1867 (2015).

23. Valdes, A. M. et al. Genetic variation in the SMAD3 gene is associated with hip and knee osteoarthritis. Arthritis Rheum. 62, 2347–2352 (2010).

24. Rodriguez-Fontenla, C. et al. Assessment of osteoarthritis candidate genes in a meta-analysis of nine genome-wide association studies. Arthritis Rheumatol 66, 940–949 (2014).

25. Casalone, E. et al. A novel variant in GLIS3 is associated with osteoarthritis. Ann. Rheum. Dis. 77, 620–623 (2018).

26. Zengini, E. et al. Genome-wide analyses using UK Biobank data provide insights into the genetic architecture of osteoarthritis. Nat. Genet. 50, 549–558 (2018).

27. Cotney, J. et al. The evolution of lineage-specific regulatory activities in the human embryonic limb. Cell 154, 185–196 (2013).

28. Herlofsen, S. R. et al. Genome-wide map of quantified epigenetic changes during in vitro chondrogenic differentiation of primary human mesenchymal stem cells. BMC Genomics 14, 105 (2013).

29. Raine, E. V. A., Wreglesworth, N., Dodd, A. W., Reynard, L. N. & Loughlin, J. Gene expression analysis reveals HBP1 as a key target for the osteoarthritis susceptibility locus that maps to chromosome 7q22. Ann. Rheum. Dis. 71, 2020–2027 (2012).

30. Luyten, F. P., Tylzanowski, P. & Lories, R. J. Wnt signaling and osteoarthritis. Bone 44, 522–527 (2009).

31. Berasi, S. P., Xiu, M., Yee, A. S. & Paulson, K. E. HBP1 repression of the p47phox gene: cell cycle regulation via the NADPH oxidase. Mol. Cell. Biol. 24, 3011–3024 (2004).

32. Scott, J. L. et al. Superoxide dismutase downregulation in osteoarthritis progression and end-stage disease. Ann. Rheum. Dis. 69, 1502–1510 (2010).

33. Liu, Y. et al. Chromatin accessibility landscape of articular knee cartilage reveals aberrant enhancer regulation in osteoarthritis. bioRxiv 274043 (2018). DOI:10.1101/274043

34. Thierry-Mieg, D., Thierry-Mieg, J. & NCBI/NLM/NIH. AceView: Gene:HBP1, a comprehensive annotation of human, mouse and worm genes with mRNAs or ESTsAceView. Available at: https://www.ncbi.nlm.nih.gov/IEB/Research/Acembly/av.cgi?db=36a&c=Gene&l=HBP1. (Accessed: 9th June 2018)

35. Gee, F., Rushton, M. D., Loughlin, J. & Reynard, L. N. Correlation of the osteoarthritis susceptibility variants that map to chromosome 20q13 with an expression quantitative trait locus operating on NCOA3 and with functional variation at the polymorphism rs116855380. Arthritis & Rheumatology 67, 2923–2932 (2015).

36. Shepherd, C. et al. Functional characterisation of the osteoarthritis genetic risk residing at ALDH1A2 identifies rs12915901 as a key target variant. Arthritis Rheumatol (2018). DOI:10.1002/art.40545

37. Wang, X., Hou, J., Quedenau, C. & Chen, W. Pervasive isoform_Jspecific translational regulation via alternative transcription start sites in mammals. Mol. Syst. Biol. 12, 875 (2016).

38. Reyes, A. & Huber, W. Alternative start and termination sites of transcription drive most transcript isoform differences across human tissues. Nucleic Acids Res. 46, 582–592 (2018).

39. Li, H. Aligning sequence reads, clone sequences and assembly contigs with BWA-MEM. arXiv [q-bio.GN] (2013).

40. Southam, L. et al. An SNP in the 5’-UTR of GDF5 is associated with osteoarthritis susceptibility in Europeans and with in vivo differences in allelic expression in articular cartilage. Hum. Mol. Genet. 16, 2226–2232 (2007).

41. Egli, R. J. et al. Functional analysis of the osteoarthritis susceptibility-associated GDF5 regulatory polymorphism. Arthritis Rheum. 60, 2055–2064 (2009).

